# Optimised genome editing for precise DNA insertion and substitution using Prime Editors in zebrafish

**DOI:** 10.1101/2025.04.23.650248

**Authors:** Yosuke Ono, Martin Peterka, Michael Love, Amir Khan, Felix Bowers, Ashish Bhandari, Euan Gordon, Jonathan S. Ball, Chrissy L. Hammond, Charles R. Tyler, Steve Rees, Mohammad Bohlooly-Y, Marcello Maresca, Steffen Scholpp

## Abstract

CRISPR/Cas9-mediated genome editing has rapidly become a popular tool for studying gene functions and generating genetically modified organisms. However, using this system, stochastic integration of random insertions and deletions restricts precise genome manipulation. Advanced CRISPR/Cas9 technologies using Prime Editors (PEs), Cas9 proteins fused with reverse transcriptase, enable programmed integration of short DNA modifications into the genome. However, its application in precise genome editing in animal models is challenging. Here, we utilise a nickase-and a nuclease-based PE to perform programmed short DNA substitution and insertion in various loci in the zebrafish genome. Whereas nickase-based PE2 mediated a higher ratio of precise prime edits to the total edits, nuclease-based PEn was more efficient for short DNA modifications, achieving up to 27.3% precise insertion. To further evaluate our approach, we inserted a nuclear localisation signal into a reporter transgene to incorporate longer fragments by prime editing. These gene modifications were transmitted to the next generation. We show that PE-mediated prime editing can efficiently manipulate genome information in zebrafish without using exogenous donor DNA.

Prime editing, CRISPR/Cas9, zebrafish, PE2, PEn

## Introduction

Genetically modified animals serve as powerful tools for studying ontogenesis and disease mechanisms in complex living systems. The zebrafish, *Danio rerio,* is a well-established model vertebrate species in cell and developmental biology due to attributes including its high fecundity in laboratory environments, transparency of the embryos, and amenability to the application of tools for transgenesis^1–3^. Zebrafish have also emerged as a powerful vertebrate model organism for studying human diseases, as they exhibit a substantial degree of genetic similarity with humans (around 70% of human genes having a corresponding gene in zebrafish)^4^. Several technologies for targeted mutagenesis, such as zinc finger nuclease (ZFN) and transcription activator-like effector nuclease (TALEN), have been employed in zebrafish to establish genetic loss-of-function mutants^5–8^. Most recently, CRISPR/Cas9-mediated genome editing has been introduced, significantly advancing target mutagenesis in nearly every model system, including zebrafish^9–13^. The CRISPR/Cas9 system has the potential to mimic disease phenotypes more accurately, allowing for the creation of identical genetic alterations corresponding to disease-causing mutations in humans. However, it has the disadvantage of stochastically generating insertions and deletions (so-called indels). This is because this technology relies on the DNA double-strand break (DSB) induced by the endonuclease at the target sequence in the genome and subsequent DNA repair. To allow precise control of genome editing, homology-directed repair (HDR)-mediated knock-in approaches using exogenous donor DNA have been utilised in zebrafish^14–16^. However, HDR-mediated precise genome editing occurs less efficiently compared with random mutagenesis.

Due to the need for precise genome editing with high efficiency, new strategies have been employed and applied in model organisms. For example, CRISPR/Cas9-mediated precise editing using Prime Editor has recently been developed, which does not rely on exogenous DNA donors. The original Prime Editor is a fusion protein of *Sp*Cas9 (H840A) nickase and Moloney murine leukaemia virus (MMLV) reverse transcriptase^17^. By combining the Prime Editor with a prime editing guide RNA (pegRNA), the resulting Prime Editor ribonucleoprotein (RNP) complex generates a single-strand break (SSB) or nick at the target site in the genome. The reverse transcriptase domain elongates an additional DNA sequence at the 3’ end of cleaved DNA, following the reverse-transcription (RT) template sequence incorporated in the pegRNA. The additional short DNA fragment that contains the intended edit is integrated into the genomic DNA through the ligation of the 3’ flap and the excision of the competing 5’ flap from the original DNA sequence^17^. The Cas9-nickase-based Prime Editor enables programmed short DNA modification (insertion, deletion and substitution), as has been shown in cultured cells, mouse and zebrafish embryos^17–20^. An alternative to the Cas9-nickase-based Prime Editor is prime editing with a Cas9-nuclease-based Prime Editor (PEn)^21–23^. PEn facilitates programmed short DNA modification through DNA repair based on homology annealing and non-homologous end joining (NHEJ) after DSB^22^. However, evidence for the successful use of Prime Editors in creating genetically modified animals is still limited. This is due to various technical challenges, including high variability in editing modes, low editing efficiency, and different delivery methods for prime editing components in living animals.

Here, we assess PE2, a nickase-based Prime Editor with additional mutations to optimise RT performance^17^, and PEn-mediated prime editing for establishing genetically modified zebrafish for programmed DNA modification in their genome. We tested nucleotide substitution and the insertion of different lengths of DNA sequences. We performed morphological assessments of these genetically modified embryos. We show that PE2 mediates prime editing more accurately for short DNA insertion and base pair substitution in zebrafish embryos, whereas editing with PEn can more efficiently insert nucleotides up to 46 bp for modulating protein function and behaviour, all without using exogenous donor DNA. We further show that stable genetically modified zebrafish lines can be generated that inherit these programmed short DNA modifications.

## Results

### Prime editing in nucleotide substitution

First, we tested the use of Prime Editors for substituting specific base pairs in the zebrafish genome. To do this, we focused on the Cereblon (CRBN) gene, which has been associated with developing resistance to thalidomide-based treatments, particularly in conditions like multiple myeloma^24,25^. Here, we first compared the functioning of nickase-based PE2 and nuclease-based PEn, in which the H840A mutation is reverted to wild type^22^, in nucleotide substitution in the zebrafish *crbn* gene (Fig. 1a), targeting the sequence encoding the amino acid residue I378 associated with the sensitivity to thalidomide and related drugs^26,27^. We designed a pegRNA to substitute 2 single nucleotides in the target sequence: one for introducing missense mutation in the I378 site (+10 A to G) and the other (+3 G to C) to prevent interaction between complementary sequences in the 5’ spacer sequence and 3’ primer binding site (PBS) and RT template sequence which potentially leads pegRNA misfolding^28,29^ (Fig. 1b). A mixture of Prime Editor proteins and chemically synthesised pegRNA was injected into zebrafish embryos at the early 1-cell stage, after which the embryos were incubated at 32°C. Genomic DNA was then extracted from these embryos at 96 hours post-fertilisation (hpf) to analyse genome edits at the target site. Amplicon sequencing of the target region showed that both PEn and PE2 integrated the prime editing substitution into the genome (Fig. 1c). We found that PE2 had a higher efficiency in precise substitutions (8.4%) compared to PEn (4.4%, Fig. 1d). Furthermore, PEn induced a higher amount of indels (Fig. 1e). The precision score, defined as the ratio of precise prime edits to the total edits, including imprecise prime edits and indels, was significantly higher with PE2 (40.8%) compared to PEn (11.4%, Supplementary Fig. 1). We also tested the refolding procedure of pegRNA by heat denaturation^28^. Refolding of the *crbn* pegRNA did not enhance prime editing substitution with both PEn and PE2 (Fig. 1d). At the *crbn* locus, PE2 is more effective for very short nucleotide substitutions compared with PEn in zebrafish.

**Fig. 1.**
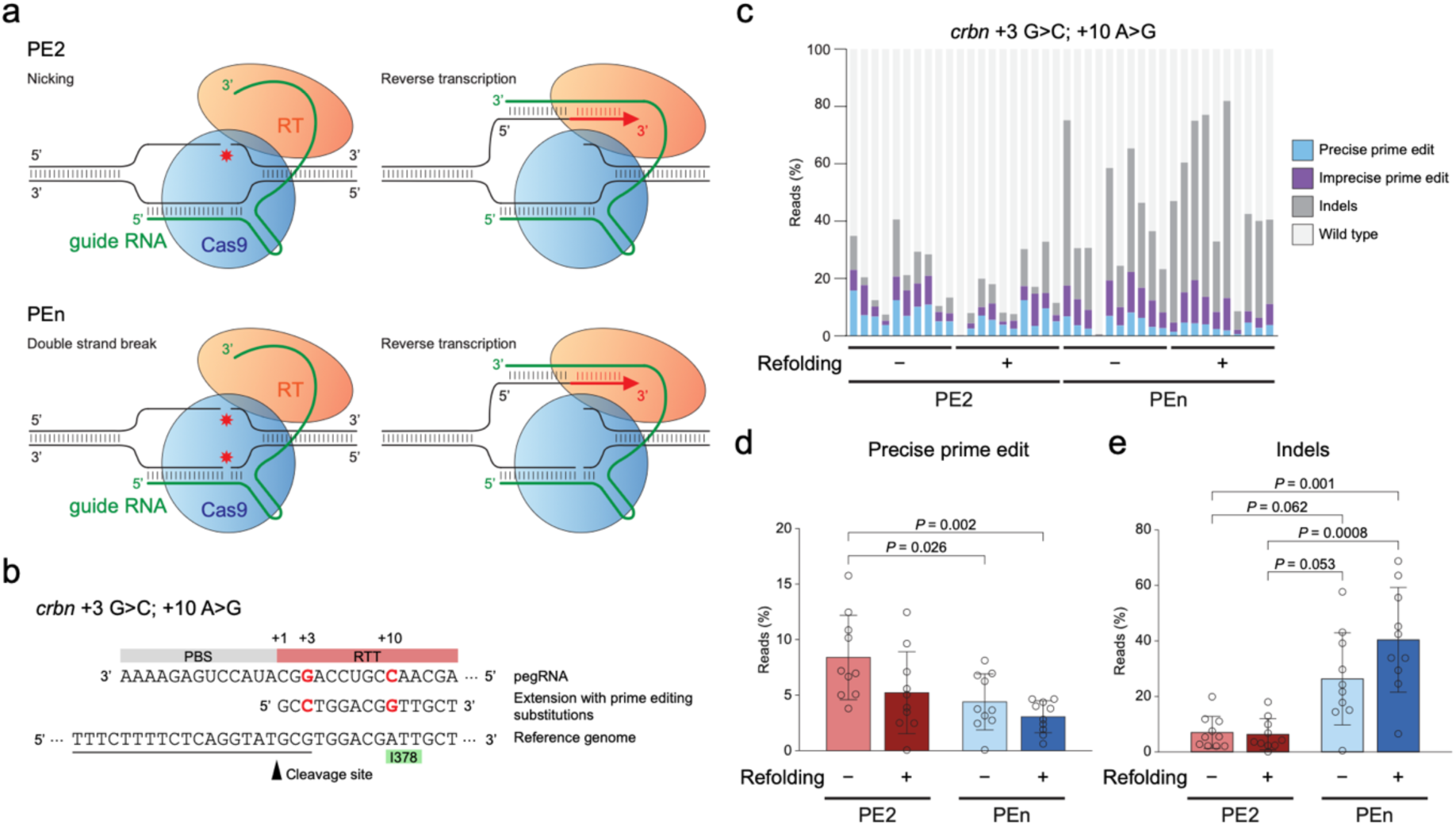
Prime editing substitution in the zebrafish *crbn* gene, comparing Cas9-nickase-based and nuclease-based Prime Editors. a, Schematic illustration of the functioning of prime editing by the Cas9-nickase-based Prime Editor (PE2) and the nuclease-based Prime Editor (PEn). b, Schematic illustration of the strategy for prime editing substitution in *crbn* gene. The nucleotides for substitution are indicated in red. The guide RNA (gRNA) target sequence is underlined. c–e, Comparison between PEn and PE2 in prime editing substitution in the *crbn* gene with the pegRNA refolding procedure. Proportions of editing outcomes in individual injected embryos are shown by amplicon sequencing (c), alongside quantitative analyses of precise prime edits (d) and indels (e) comparing experimental conditions (n = 10 per group). P-values were determined using one-way ANOVA with Tukey’s multiple comparison test in (d), and the Kruskal-Wallis test with Dunn’s multiple comparison test in (e). Error bars in the bar graphs represent the mean and standard deviation, and individual data points indicate values from single injected embryos.

### Prime editing insertion of 3 bp DNA fragment

Next, we tested the functioning of the Prime Editors in the precise insertion of a 3 bp stop codon into the coding sequence of the endogenous target gene to generate a mutant allele that produces a precisely truncated protein. As a proof-of-concept, we chose the cognate receptor gene *ror2*. Genetic mutations in the *ROR2* gene in humans can cause the autosomal recessive Robinow syndrome, leading to short stature, distinctive facial features, and skeletal abnormalities, including short limbs and a curved spine (*scoliosis*)^30–33^. In zebrafish, the inhibition of Ror2 signalling function affects the convergence and extension of axial cells during gastrulation and the elongation of the embryo body, leading to a broader and shorter body axis^34–37^, as well as the patterning of cranial tissues^38^, and thus would be an excellent model for studying many aspects of the Robinow syndrome.

Sequence comparison suggested that the zebrafish W722X allele corresponds to the human disease-related W720X mutation generating a premature stop codon (TGA) in the tyrosine kinase domain (Fig. 2a). Therefore, we aimed to establish the zebrafish Robinow W722X model by designing pegRNA and single primed insertion gRNA (springRNA)^22^ generating a similar stop codon into the sequence of exon 9 in zebrafish *ror2.* The pegRNA contained a 3 nt RT template sequence for the stop codon to be integrated, 13 nt PBS sequence, and 13 nt sequence to extend the homology arm for the DNA integration via homology annealing (Fig. 2b and c). We compared this strategy to using a springRNA combined with PEn, without the template sequence for the homology arm, to insert the stop codon via NHEJ (Fig. 2b and c). We microinjected combinations of Prime Editor mRNA and guide RNA, that is, PE2/pegRNA, PEn/pegRNA and PEn/springRNA at the one-cell stage in zebrafish embryos. The sequence in *ror2* exon 9 was amplified by PCR from a pool of genomic DNA obtained from 10 injected embryos, and the induction of DNA modification was assessed by a T7 endonuclease I (T7E1) assay. We observed the cleavage of PCR products in the samples injected with PEn/pegRNA and PEn/springRNA combinations, indicating sequence modification in the target site (Fig. 2d). We were unable to detect obvious cleavage of heteroduplex DNA with PE2/pegRNA combination, suggesting that PE2 induced fewer editing events compared to PEn.

**Fig. 2.**
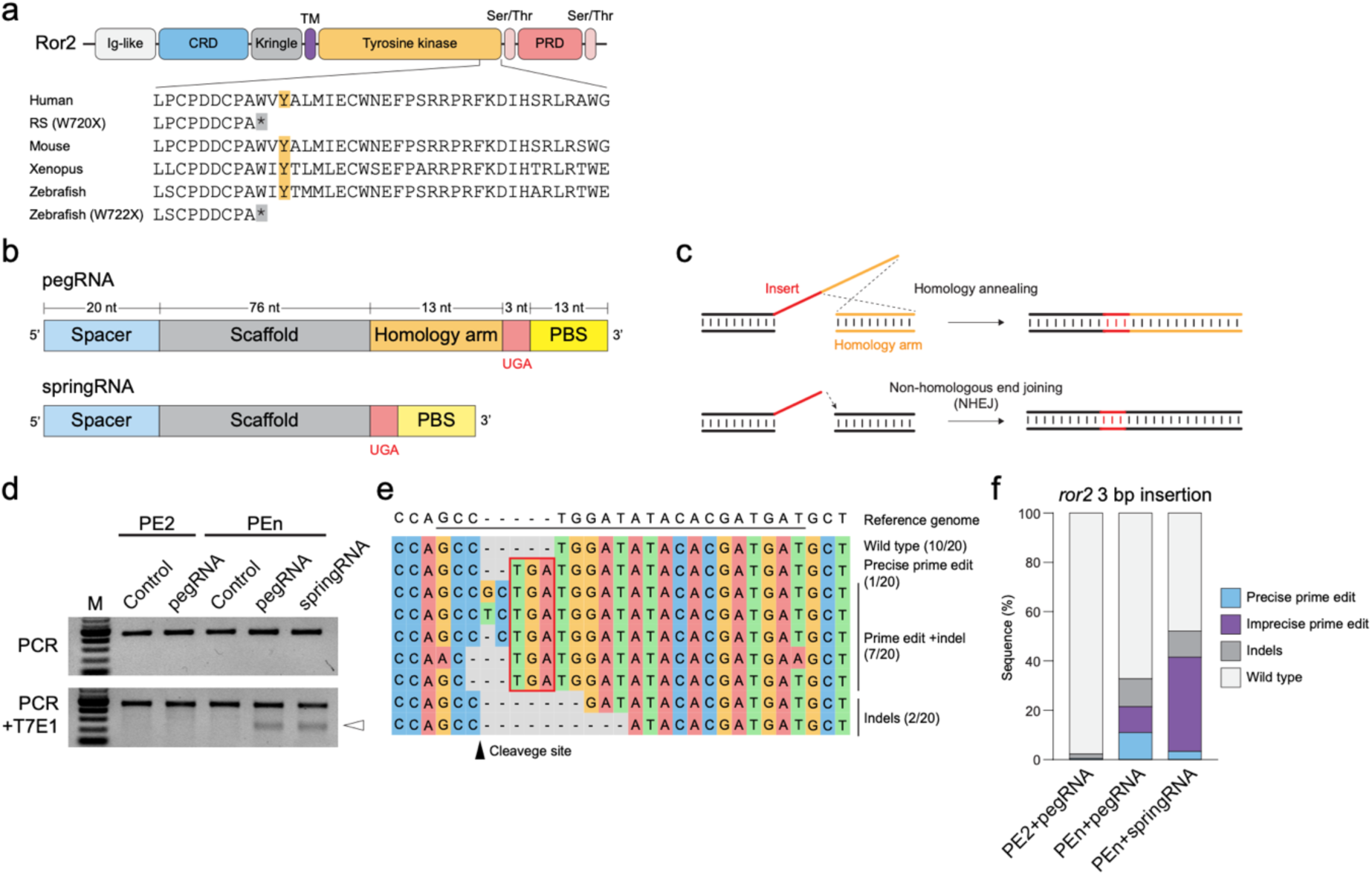
Prime editing insertion in zebrafish *ror2* gene using Cas9-nuclease-based Prime Editor. a, Schematic illustration of the functional domains in the Ror2 protein and alignment of partial amino acid sequences within the tyrosine kinase domain. Sequences from multiple species, including those related to Robinow syndrome (RS) in humans W720X and the corresponding zebrafish W722X mutant, are aligned. The conserved tyrosine residue is highlighted. b, Schematic illustration of guide RNA (gRNA) designs for prime editing insertion in *ror2*. c, Schematic illustration of prime editing insertion by Cas9-nuclease-based Prime Editor (PEn). An additional DNA fragment, reverse-transcribed at the target cleavage site, containing the programmed insertion, is integrated into the genome via homology-directed repair or non-homologous end joining. d, Agarose gel images of genomic PCR products from embryos injected with Prime Editor mRNA and pegRNA/springRNA. PCR products of the *ror2* target region (top) and those after digestion with T7 endonuclease I (T7E1, bottom). e, Sequence alignment of the edits in the *ror2* target site obtained from embryos injected with PEn/springRNA. Prime editing insertion (TGA) is outlined, and the gRNA target sequence is underlined. f, Quantitative comparison of editing outcomes using different combinations of Prime Editor and gRNA. The proportion of sequence reads with each type of edit in amplicon sequencing is presented.

The PCR products were also cloned into a cloning vector, and the target sequences in randomly selected clones were analysed. Various types of edits in the target site were observed in the clones from the PEn injected samples, including precise insertion of the stop codon and/or random indels (Fig. 2e), although no sequence edits in the target site were identified from the PE2 injected sample (0/44 clones; Supplementary Fig. 2). Subsequently, we performed amplicon sequencing to quantify the proportion of each genome edit. We found that the proportion of precise prime editing (only TGA insertion without any indels) was higher when using PEn/pegRNA combination (10.3%) compared to PEn/springRNA (4.0%) and PE2/pegRNA (0.4%, Fig. 2f). In addition, springRNA showed a higher proportion of imprecise prime edit (TGA insertion with random indels; 37.5%) than pegRNA (11.1% and 0.2% with PEn and PE2, respectively; Fig. 2f). The precision scores for the PE2/pegRNA, PEn/pegRNA, and PEn/springRNA combinations were 58.9%, 32.6%, and 7.8%, respectively. This suggests that prime editing via homology annealing with pegRNA was more accurate than via NHEJ using springRNA.

### Comparison of the mode of Prime Editor delivery as mRNA or RNP complex

Next, we compared PEn and PE2 in combination with different strategies of Prime Editor delivery. Here, we performed amplicon sequencing with genomic DNAs extracted from individual embryos injected with Prime Editor mRNA or RNP complex with the *ror2* pegRNA (Fig. 3a). We found that the proportion of precise prime edit is significantly increased with PE2 RNP complex (11.5%) compared to PE2 mRNA (1.3%), whereas the there is no significant difference between PEn RNP complex (20.5%) compared to PEn mRNA (18.8%; Fig. 3b). However, with both delivery methods of the Prime Editors, the proportion of precise prime edit was significantly higher with PEn compared to PE2, supporting our previous results (Fig. 2f and 3b) although, PEn-mediated prime editing insertion outperformed HDR-mediated knock-in using single-stranded donor DNAs (2.3% and 5.2%; Fig. 3c). Furthermore, PEn also integrated a notably higher number of indels that are unrelated to prime editing (Fig. 3d), probably due to the DNA DSB induced by its nuclease activity as the indel frequency was similar level to the conventional HDR-mediated knock-in using Cas9 RNP complex (Supplementary Fig. 3a). These resulted in lower precision scores with PEn (34.3% and 28.2% for mRNA and RNP delivery, respectively) compared to PE2 (71.6% and 43.4% for mRNA and RNP delivery, respectively; Supplementary Fig. 3b). We also tested the generation of *ror2* W722X allele by single nucleotide substitution (TGG to TGA). Notably, PEn-mediated prime editing introduced targeted substitution (3.7%) more efficiently than PE2 (2.0%) in *ror2*, whereas PEn induced a higher amount of indels resulting in a lower precision score (14.8%) compared to PE2 (25.4%; Supplementary Fig. 3c–e). These findings suggest that the PEn-induced DNA DSBs are more likely to undergo repair events (both precise and imprecise) compared to the nickase-based PE2 method. Although PEn and PE2 showed distinct editing outcomes, both Prime Editors did not show a significant difference in the frequency of non-specific editing at 3 potential off-target sites in the genome, and their specificity was comparable to HDR-mediated knock-in using donor-DNA into the same target site (Supplementary Fig. 3f and g).

**Fig. 3.**
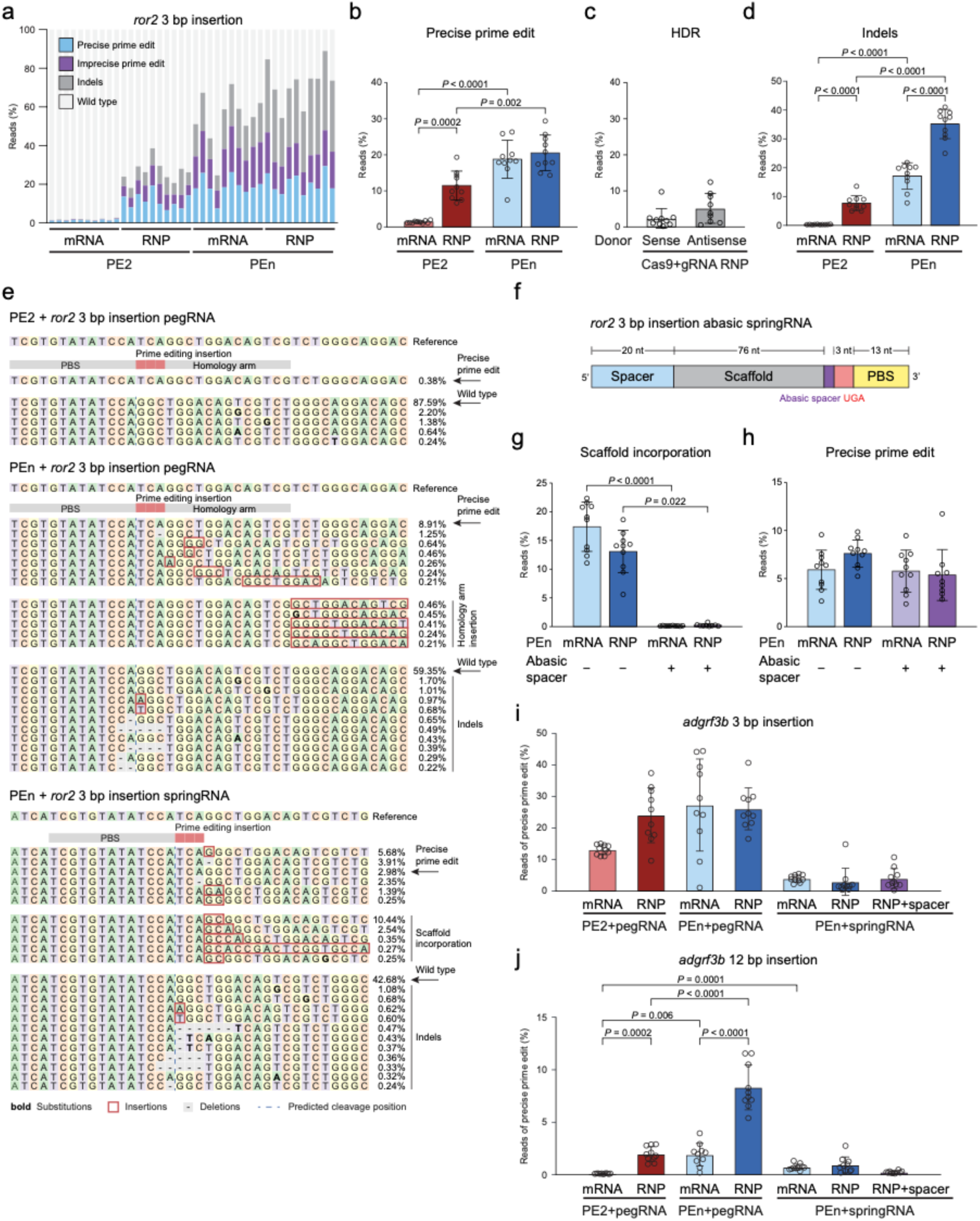
Comparing different prime editing approaches in zebrafish embryos. a and b, Comparison between PEn and PE2 in prime editing insertion into *ror2* with various delivery methods. The proportion of editing outcomes in individual injected embryos was assessed through amplicon sequencing (a), along with a quantitative analysis of precise prime edit (b) and indels (d) across experimental conditions (n = 10 per group). c, Proportion of reads with precise HDR-mediated insertion in *ror2* using single-stranded sense or antisense donor DNAs with 40 bp homology arms. d, Proportion of reads with indels comparing PEn and PE2 in prime editing insertion into *ror2*. e, Sequence alignment of edits in prime editing insertion in *ror2* via PE2 mRNA/pegRNA (top), PEn mRNA/pegRNA (middle), and PEn mRNA/springRNA (bottom) combinations. f, Schematic illustration of springRNA design featuring an abasic RNA spacer for prime editing insertion in *ror2*. g and h, Proportion of reads with scaffold incorporation (g) and precise prime edit (h) as determined by amplicon sequencing of prime editing insertion in *ror2* using control and abasic springRNA (n = 10 per group; one sample of the PEn RNP/abasic springRNA combination was excluded from analysis due to low read count). i and j, Prime editing insertion in the *adgrf3b* gene using PE2 and PEn. The proportion of reads with a precise 3 bp (i) and 12 bp insertion (j) was evaluated (n = 10 per group). P-values were determined by Welch’s one-way ANOVA with Dunnett T3 multiple comparison test in b, d, and j, and Kruskal-Wallis test with Dunn’s multiple comparison test in g. Error bars in bar graphs represent the mean and standard deviation, and each individual data point indicates the value from a single injected embryo.

### Inclusion of abasic spacer into springRNA to prevent guide RNA scaffold insertion

In the analysis of the sequence edits in the *ror2* gene by PEn-mediated prime editing, we found that a substantial proportion of the amplicons derived from embryos injected with the PEn/springRNA combination contained imprecise prime edits with additional nucleotides that correspond to the sequence of guide RNA scaffold adjacent to the RT template (Fig. 3e), as reported previously^18^. This finding suggests that the readthrough of RT from the template sequence into the guide RNA scaffold causes unintended insertions, as has also been reported in human cell lines^17,39^. Aiming to control the termination of RT in prime editing, we synthesised a modified springRNA in which an abasic RNA spacer was inserted between the RT template and guide RNA scaffold (Fig. 3f). Microinjection of the abasic springRNA together with PEn mRNA or RNP complex significantly reduced the unintended insertion of guide RNA scaffold sequence into the target site (Fig. 3g). However, the proportion of precise prime edits did not show a significant improvement (Fig. 3h). Due to the significantly reduced number of the imprecise prime edits, the precision scores increased from 9.6% to 21.7% and from 10.6% to 14.2% with PEn mRNA and RNP complex, respectively, through the blocking of scaffold sequence incorporation (Supplementary Fig. 3h). These results suggest that the read-through process is independent of precise genome editing events in zebrafish embryos, unlike in human cultured cells^39^.

### Prime editing in adgrf3b gene using PEn

Next, we tested the PEn-mediated prime editing targeting at a different locus, here the *adgrf3b* gene. A previously published study has reported that the efficiency of prime editing in the *adgrf3b* gene using PE2 RNP complex was decreased from 18.0% to 0.1% by extending the length of prime editing insertion from 3 bp to 12 bp^18^. Therefore, we compared the efficiency of our nuclease-based PEn to PE2 by targeting the same sequence in the *adgrf3b* gene to integrate the 3 bp or 12 bp insertion. Both PEn/pegRNA and PEn/springRNA combinations successfully integrated the programmed 3 bp and 12 bp insertion into the target site in the injected embryos. Here, the mean proportion of precise prime edit for 3 bp insertion using PEn mRNA with pegRNA and springRNA was 27.3% and 4.0%, respectively, and 26.1% and 2.9% using PEn RNP complex (Fig. 3i). For 12 bp insertion, the mean proportion of precise prime edit using PEn mRNA with pegRNA and springRNA was and 1.9% and 0.7%, respectively, and 8.3% and 0.9% using PEn RNP complex (Fig. 3j). The precision scores of PEn mRNA with pegRNA and springRNA were 42.1% and 4.6% for a 3 bp insertion and 8.6% and 1.1% for a 12 bp insertion, respectively (Supplementary Fig. 3i), suggesting that both the efficiency and accuracy of prime editing declined with an increase in insertion length. Notably, the PEn RNP complex with pegRNA performed more efficiently with the 12 bp insertion compared to the PE2 system, this suggests that PEn-generated DNA DSB accepts longer insertion than a nickase-based Prime Editor in zebrafish embryos. Inclusion of an RNA or DNA spacer into the springRNAs did not improve prime editing insertion in *adgrf3b*, whereas the incorporation of guide RNA scaffold was significantly reduced similarly to the *ror2* locus (Supplementary Fig. 3j and k).

### Establishing stable ror2^W722X^ mutant zebrafish

Having demonstrated the programmed insertion via PEn-mediated prime editing in injected embryos, we wanted to establish if these precise edits are passed on to the next generation, as this is a prerequisite for establishing a disease model. We, therefore, aimed to develop stable *ror2*^W722X^ mutant zebrafish. Therefore, first, we injected embryos with PEn RNP complex and the *ror2* springRNA and raised the embryos to adults, and then we outcrossed them with wild-type fish to collect F1 embryos. Through genotyping 16 embryos from each F0 adult fish, we obtained F1 embryos harbouring the *ror2*^W722X^ mutation. Six out of 10 F0 adults were founders of the allele, with the rate of germline mosaicism varying between 6.3% (1/16) and 31.3% (5/16; Fig. 4a). We raised the F1 heterozygous *ror2*^W722X^ fish and inbred them to generate homozygous *ror2*^W722X^ mutant embryos in the F2 generation. The morphology of the zygotic *ror2*^W722X^ mutant embryo and larvae was similar to wild-type siblings, and the mutant animals were both viable and fertile (Fig. 4b, c and e). Subsequently, we generated maternal-zygotic (MZ) mutants by in-crossing homozygous *ror2*^W722X^ male and female fish. The MZ *ror2*^W722X^ mutant larvae at 5 dpf showed a mild curvature of the tail tip and shorter body length (Fig. 4d and f), which are similar phenotypes to the MZ mutant larvae of the loss-of-function alleles in *ror2* reported previously^34,38^. Zygotic and MZ mutant adults lacked craniofacial sensory organs, nasal and maxillary barbels (Fig. 4g and h, and Supplementary Fig. 4), and 1 year old mutant showed altered jaw morphology with a significantly altered aspect ratios in the mandible, leading to a less protrusive lower jaw (Fig. 4i–m), phenocopying the larval and adult phenotypes previously described for zebrafish *ror2* mutants^38^. These phenotypes also resemble the craniofacial defects in *Ror2* mutant mice^40^ and the micrognathia observed in human patients with Robinow syndrome^32,33^. This suggests that the human disease-related *ror2*^W722X^ mutation in the tyrosine kinase domain affected the Wnt/PCP signalling similarly to the observation in patients with Robinow syndrome. This set of data suggests that PEn-based editing can be used to establish stable genetically modified fish lines for disease modelling.

**Fig. 4.**
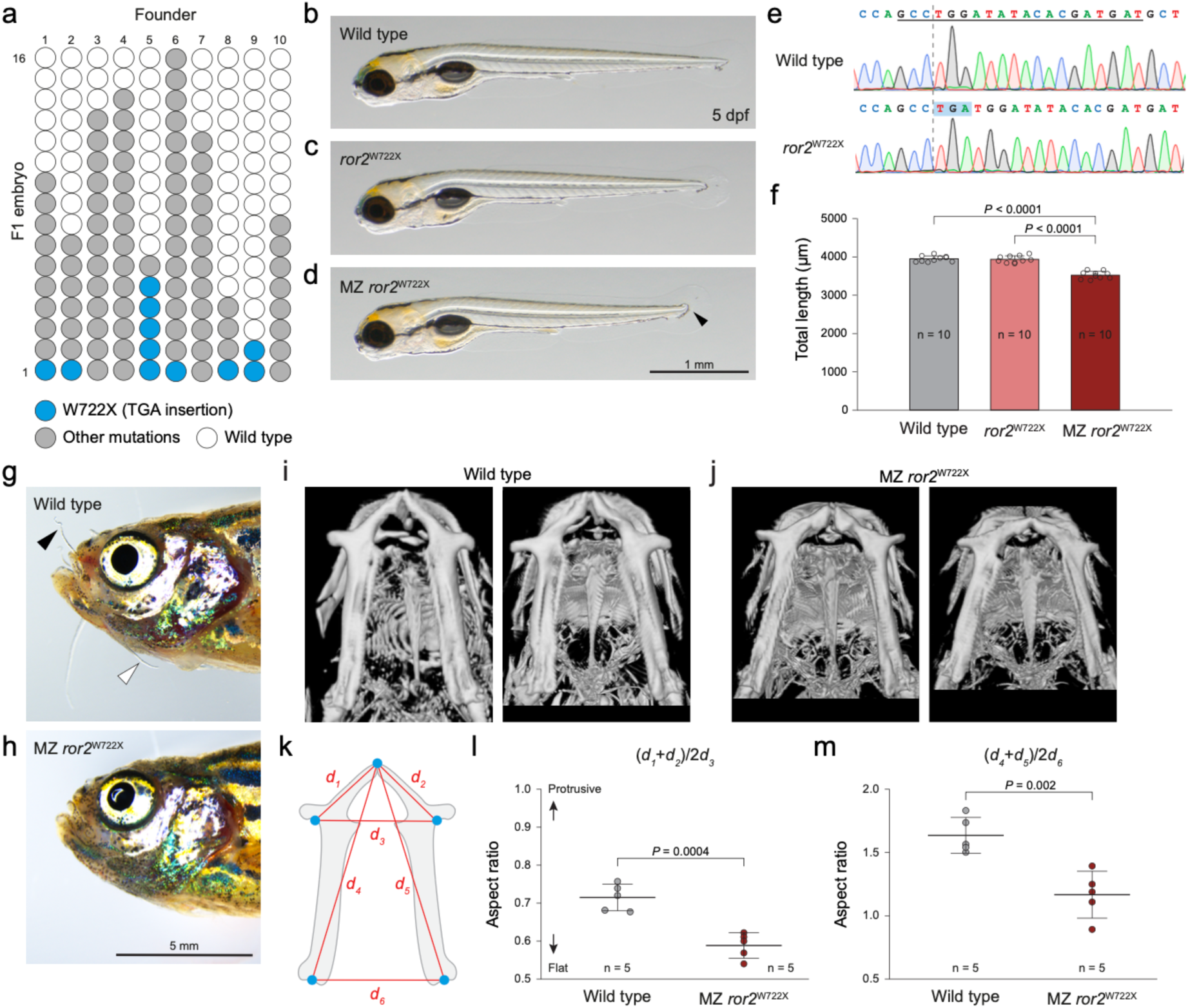
Generation and characterisation of stable *ror2^W722X^* mutant. a, Summary of the screening of F1 embryos. Sixteen embryos were genotyped per F0 founder. Each circle represents the genotype of a single embryo. The embryos were obtained by outcrossing the injected founder with wild-type fish; thus, all embryos with the mutation are heterozygous. b-d, Lateral images of wild type (b), zygotic *ror2^W722X^* mutant (c), and maternal-zygotic (MZ) *ror2^W722X^*mutant (d) larvae at 5 days post-fertilisation (dpf). e, Sanger sequencing chromatogram of wild type (top) and *ror2^W722X^* mutant (bottom) at the prime editing target site in *ror2*. Prime editing insertion (TGA) is highlighted. The target sequence of the guide RNA is underlined, and the cleavage site of Cas9 is indicated by a dotted line. f, Quantitative analysis of total length comparing wild type, zygotic *ror2^W722X^* mutant, and MZ *ror2^W722X^*mutant at 5 dpf. Each data point on the graph represents the value from a single larva (n = 10 per group). P-values were determined by one-way ANOVA with a Tukey multiple comparisons test. Error bars represent the mean and standard deviation. g and h, Lateral images of 1 year old wild type (g) and MZ *ror2^W722X^* mutant (h). Nasal and maxillary barbels are indicated by black and white arrowheads, respectively. i and j, Reconstructed computed tomography images of cranioskeletal morphologies in 1 year old wild type (i) and MZ *ror2^W722X^* mutant (j). The images are ventral view and anterior is to the top. k, Schematic illustration of mandibular bone from the ventral view and the location of geographical points for annotation to measure linear distances and calculate the aspect ratios. l and m, Quantitative analysis of the aspect ratios in the mandible comparing wild type and MZ *ror2^W722X^*mutant. Aspect ratio of anterior part (l) and whole structure (m) of the mandible was analysed. The p-values were determined using unpaired t-test. Error bars represent the mean and standard deviation, while individual data points on the graph indicate values from single reconstructed image.

### Insertion of nuclear localisation signal into transgenic reporter gene

After the successful insertion of the 3 bp stop codon into the *ror2* gene to establish a zebrafish model of the Robinow syndrome and successful 3 and 12 bp insertions into the *adgrf3b* gene, we tested whether PEn-mediated prime editing can be used to integrate longer insertions. Since we anticipated that these events would be rare, we employed a strategy that enabled us to screen for prime edit events and distinguish between imprecise prime edits and random indels at the cellular level in living zebrafish embryos, all without the need for time-consuming sorting and genetic analysis. This was achieved using an eGFP-transgenic fish line with an introduced nuclear localisation sequence (NLS) of 27 bp using PEn. We anticipated three phenotypes: cytosolic GFP localisation for unsuccessful editing, no GFP expression for imprecise edits, and nuclear GFP expression for precise prime editing insertion (Fig. 5a). We subsequently mapped these effects in individual cells using high-resolution confocal microscopy in living zebrafish larvae. Specifically, we tested the insertion of the NLS sequence into the *smyhc1:gfp* transgene by prime editing. The *smyhc1:gfp* reporter gene is expressed explicitly in the superficial slow-twitch muscle fibres in the body trunk (Supplementary Fig. 5a), which are mononuclear during embryonic development^41^. We designed pegRNAs that target the N-terminal end of the eGFP coding sequence in the *smyhc1:gfp*^i104^ transgenic allele and contain RT template sequence for inserting c-myc NLS^42–44^. We expected that precise integration of the 30 bp sequence, including 27 bp c-myc NLS and 3 bp spacer, translocates the eGFP expression from the cytoplasm to the nucleus in the muscle fibres, whereas reporter expression is lost in the cells with imprecise prime edits or random indels by frameshifting in the eGFP coding sequence (Fig. 5a). The embryos injected with PEn RNP complex showed mosaic expression of eGFP in the cytoplasm of muscle fibres, lack of GFP expression in some fibres along with the accumulation of eGFP signals in a few cell nuclei (Fig. 5b). Amplicon sequencing of the target region in the genome showed that the average efficiency of precise prime editing insertion in the whole injected embryo was 8.5% with a precision score of 9.7% (Fig. 5c). We also compared the prime editing efficiency to that using a longer pegRNA in which PBS sequence and template sequence of the homology arms were extended from 10 nt to 13 nt and 14 nt to 36 nt, respectively. However, the prime editing efficiency of the longer pegRNA was significantly decreased, possibly due to increased RNA complexity leading to altered secondary structures (Fig. 5c).

**Fig. 5.**
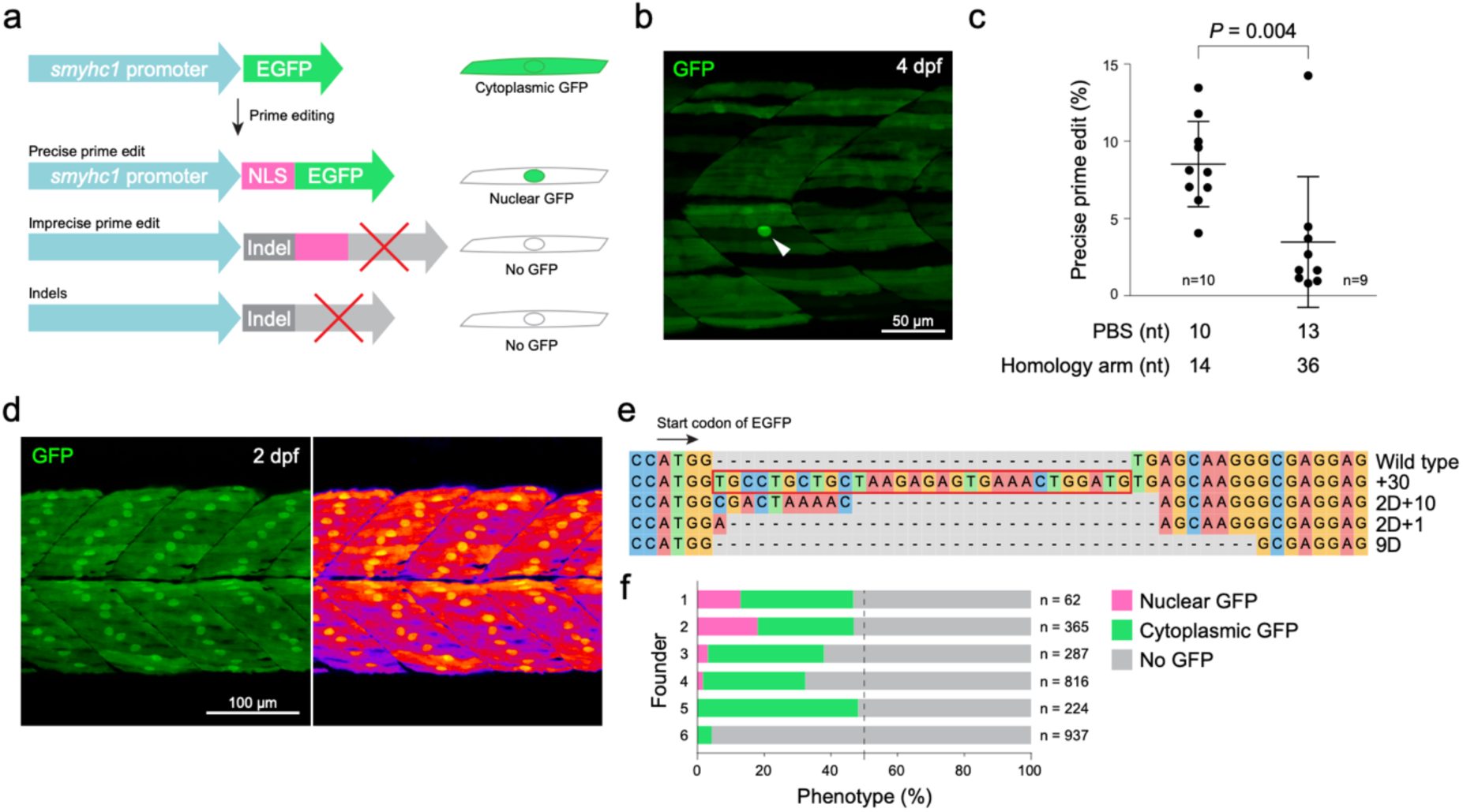
Prime editing to insert a nuclear localisation signal sequence into the *smyhc1:gfp* transgene. a, Schematic representation of the prime editing insertion of the nuclear localisation signal (NLS) sequence into the *smyhc1:gfp* transgene and the expected eGFP expression in slow-twitch muscle fibres. b, Confocal microscopy image of the trunk muscle in *smyhc1:gfp* larvae at 4 days post-fertilisation (dpf) that were injected with the PEn RNP complex for the prime editing NLS insertion. Putative nuclear GFP expression is indicated by an arrowhead. Anterior is positioned to the left. c, Quantitative analysis of the efficiency of precise NLS insertion via amplicon sequencing. Two pegRNAs of differing lengths are compared (n = 10 per group, with one sample excluded from the analysis due to low read count). The p-value was determined using the Mann-Whitney U test. Error bars represent the mean and standard deviation, while individual data points on the graph indicate values from single injected larvae. d, Confocal microscopy images of the trunk muscle of F1 larvae exhibiting nuclear GFP expression at 2 dpf, obtained from founder 1 (see panel f). The GFP fluorescence channel (left) and pseudo-colour (right) images are shown. Anterior is to the left. e, Sequence alignment of the edits in the target site obtained from F1 embryos exhibiting nuclear GFP expression (founder 1). The NLS sequence is outlined in red. f, Summary of F1 embryo screening. F1 embryos obtained from 6 founders are categorised based on the GFP expression pattern. The embryos were produced by outcrossing the heterozygous *smyhc1:gfp* founder injected with the PEn RNP complex to wild type; thus, half of the embryos are expected to be negative for the *smyhc1:gfp* transgene.

Next, to establish stable genetically modified zebrafish lines with the NLS insertion, we raised F0 injected fish to adults, outcrossed them with wild type, and screened the F1 embryos for their GFP expression. Although 8.0% (7/87) of injected F0 fish showed mosaic eGFP expression with nuclear eGFP at 3 dpf, there was a difficulty in estimating prime editing efficiency by the observation, as muscle fibres showed variable ratios of nuclear eGFP to cytoplasmic eGFP; therefore, we did not pre-screen them for raising. We successfully isolated embryos showing pronounced eGFP expression in the nuclei of slow-twitch fibres (Fig. 5d and Supplementary Fig. 5b). We confirmed the integration of the 30 bp sequence into the eGFP coding sequence (Fig. 5e and Supplementary Fig. 5c). Notably, multiple types of edits were also identified in the amplicons obtained from the single embryo with nuclear eGFP expression, suggesting that the *smyhc1:gfp*^i104^ transgenic allele contains at least 4 copies in this locus. Each copy was targeted and edited individually by PEn/pegRNA complex (Fig. 5e and Supplementary Fig. 5c). Here, 4 F0 adult fish out of 6 screened were identified as founder fish of the prime editing insertion in the target transgene. The proportion of the embryos showing nuclear eGFP expression varied between 1.4 to 20.2% (Fig. 5f).

To challenge the length of insertion using PEn in zebrafish embryos further, we also tested the insertion of a 46 bp attP sequence for attP/attB recombination by phiC31 integrase, which could expand the application of PEn for transgenesis^45–47^. Microinjection of PEn RNP complex with springRNA that contains RT template for inserting a 46 bp attP sequence into *wls* gene achieved the programmed insertion at the efficiency of 1.4% with a precision score of 2.9%, and the efficiency was comparable to HDR-mediated knock-in into the same target site using single-stranded donor DNAs (1.4% and 0.5%; Supplementary Fig. 6). Our findings indicate that PEn-mediated precise DNA insertion is a practical tool for the accurate integration of functional motifs with a length of up to 46 bp within a target gene in zebrafish.

## Discussion

### The need for precise genome editing

Target sequence-specific mutagenesis and transgenesis are essential tools for analysing gene functions, and labelling expressed proteins, cellular components and specific cell types *in vivo*. Induction of DSB in the genome DNA by ZFN, TALEN and CRISPR/Cas9 technologies enable random mutagenesis and donor DNA-based knock-in in the target sequence. In addition to the conventional HDR-mediated knock-in, multiple systems for targeted integration into the genome in zebrafish have been reported^47,48^. Adding further to this, Cas9-nickase fusion proteins with cytidine or adenine deaminase, so-called Base Editors^49,50^, enable single nucleotide substitution in zebrafish to generate human disease models^51–56^. Despite advancements in gene editing technologies, however, the application of Prime Editors for the generation of genetically modified animals remains restricted^18,57–59^. This limitation can be attributed to several technical hurdles, including substantial variability in editing modes, suboptimal editing efficiency, and complications associated with the delivery of prime editing components *in vivo*.

In this study, we established programmed short DNA modification in the genome using Prime Editors and generated stable genetically modified zebrafish lines with 3 and 30 bp insertions. Firstly, we utilised PE2 for successful nucleotide substitution in a target sequence associated with drug sensitivity and PEn to generate a zebrafish model for human genetic disease. Specifically, we established a stable *ror2*^W722X^ mutant line, which genocopies the disease-related W720X mutation in the human *ROR2* gene. Various mutations have been identified in the human *ROR2* gene as the causes of Robinow syndrome, and different types of mutation lead to distinct phenotypes and inheritance patterns, possibly due to nonsense-mediated decay of mutated mRNA^60^, the mechanism which is also essential for embryonic development and tissue homeostasis in zebrafish^61,62^. Our establishment of the *ror2*^W722X^ line supports the use of prime editing to generate zebrafish models of disease-relevant alleles in cases where short, programmed substitutions or insertions are sufficient. Using prime editing and base editing by Base Editors will open a wider window for precise genome editing in zebrafish for both drug discovery and human disease models.

Secondly, we inserted a functional motif in existing lines to control the features of a protein. Here, specifically, we inserted an NLS sequence into the *smyhc1:gfp*^i104^ transgene and assessed the prime editing *in vivo* by translocating eGFP expression. The same approach can be applied to tag endogenous proteins and control their localisation and transport. The pegRNAs we used to insert NLS were designed in the coding sequence of the N-terminal region of eGFP, and therefore, the strategy can be used for other established eGFP reporter zebrafish lines to convert their GFP expression from the cytoplasm to the cell nuclei and as a control to assess prime editing *in vivo* morphologically.

Previous studies have reported a low efficiency of prime editing insertion. In those studies it is proposed that the length negatively correlates with prime editing efficiency, while the MLH1 protein influences editing outcomes for short sequences up to 13 bp in cultured cells^63–65^. Indeed, we observed the declined efficiency and accuracy of prime editing with extended insertion in zebrafish embryos. However, the insertion of the 30 bp NLS into the *smyhc1:gfp* transgene showed similar or higher editing efficiency to the 12 bp insertion into the *adgrf3b* gene, suggesting that other factors such as spacer sequence and/or complexity of pegRNA also affect the prime editing efficiency^63^. Enhancing pegRNA stability and preventing degradation can further improve editing efficiency^28,29,66^. Finally, co-expressing effector proteins with Prime Editor to suppress undesired DNA repair pathways or protect pegRNA from degradation has been suggested to enhance editing^23,64,67^. Evaluation of these technologies in zebrafish^68,69^, as well as codon optimisation of Prime Editors and adaptation to RT at lower temperature, will advance precise genome manipulation *in vivo* further.

### The underlying mechanism of PEn-mediated precise genome editing

Here, we demonstrated that a nuclease-based PE can efficiently integrate DNA insertions between 3–46 bp in zebrafish embryos. By utilising pegRNA and springRNA, we show that the integration is based on both homology annealing and NHEJ, as suggested in cell culture^22^. Although pegRNA performed better under the tested conditions than springRNA in both efficiency and accuracy at the tested target loci, springRNA could be advantageous when longer PBS and/or RT template sequences are necessary, as it lacks a template for homology arms. We also observe that the repair outcome varies depending on the delivery mode of the PEn. Microinjection of PEn mRNA leads to fewer indels than supplying the PEn as an RNP complex, although there was no significant difference in the level of precise prime edit in the 3 bp insertion at *ror2* and *adgrf3b* loci (Fig. 3). We hypothesise that the RNP complex acts faster than the mRNA and a potential explanation could be that the relatively rapid integration of random indels is more effective than the prime editing reaction in the early zebrafish embryo (0–3 hpf), when cell cycle time is significantly shorter (approximately 25 min). However, the precise DNA repair mechanism and accessibility to a target site in the genome for the prime editing reaction in relation to the cell cycle duration need to be explored in the future.

We also show that nuclease-based PE has its limitations. For example, we found that the nickase-based PE2 showed higher precision scores in nucleotide substitution and short DNA insertion, as shown for *crbn* (Fig. 1 and Supplementary Fig. 1) and *ror2* (Supplementary Fig. 3). Further analyses will be required to compare nuclease-based and nickase-based Prime Editors, focusing on the distance between the substitution sites and Cas9 nicking/cleavage site. Furthermore, as PEn functions via DSB of genomic DNA, insertion can potentially cause off-target effects, large deletions, and chromosomal rearrangements^70–72^, whereas the level of large deletions in PEn-mediated prime editing is comparable to that with the nickase-based PE2 in cultured cells^22^. It will be interesting to see whether these adverse effects that occurred in injected zebrafish can be neutralised by outcrossing against wild-type fish in the next generations.

In conclusion, we have established genetically modified zebrafish with programmed short DNA modifications by microinjecting Prime Editor mRNA or RNP complex with gRNAs that integrate DNA modification via different DNA repairing pathways. PEn-mediated prime editing insertions were germline-transmitted to the next generation in the tested loci with measurable efficiency to establish stable genetically modified zebrafish lines. Our data establish prime editing-mediated insertion in zebrafish embryos as a practical donor DNA-free approach for precise short DNA modification, although editing efficiency and precision remain locus-and edit-dependent. The method can be employed to manipulate coding sequences in endogenous genes and to insert functional DNA motifs for protein tagging and modulating gene expression. Our data support the utility of PE2 for accuracy and PEn for efficiency in short DNA modifications in F0 injected zebrafish, while broader comparison of germline transmission efficiencies between prime editing systems will require future work.

## Materials and Methods

### Zebrafish strains and husbandry

Wild-type zebrafish strains WIK and AB and the *Tg(smyhc1:gfp)*^i104^ transgenic line^41^ were used to provide embryos. The collected embryos were raised at 28°C. Adult fish stocks were reared and maintained in the Aquatic Resources Centre at the University of Exeter under the conditions stated in the previous study^73^. The experimental procedures involving the animals were reviewed and approved by the United Kingdom Home Office under the Animals Scientific Procedures Act 1986 (project license: PP3975835) and received ethical approval from the Animal Welfare and Ethical Review Body at the University of Exeter.

### Preparation of pegRNA and springRNA

The sequences of the pegRNA and springRNA for prime editing in the *crbn*, *ror2*, and *wls* genes, and for the *smyhc1:gfp* transgene, are listed in Supplementary Table 1. The pegRNAs for prime editing insertion in the *adgrf3b* gene have been reported in a previous study^18^. The first three nucleotides and inter-nucleotide linkages at both ends of the gRNA were chemically modified with 2’-O-methylation and phosphorothioate, respectively. Chemically modified synthetic gRNAs were obtained from Integrated DNA Technologies. The springRNAs with an RNA or DNA spacer were sourced from Horizon Discovery and GenScript Biotech. Synthesised gRNAs were dissolved in nuclease-free duplex buffer (Integrated DNA Technologies) at 100 µM and stored at −20°C.

### Microinjection

Microinjection into one-cell stage embryos was carried out under a stereomicroscope using a FemtoJet 4x microinjector (Eppendorf). To prepare an injection mixture for prime editing, 1.25 µl of 600 ng/µl Prime Editor mRNA or 0.6 µl of 10 mg/ml purified Prime Editor protein was combined with 2.5 µl of 12 µM gRNA and 0.5 µl of 0.5% phenol red (Sigma), and the volume was adjusted to 5 µl with 2 M potassium chloride. PEn and PE2 mRNA were synthesised from the linearised plasmids^22^ using the mMESSAGE mMACHINE T7 kit (Invitrogen), following the manufacturer’s instructions. Purified Prime Editor proteins were prepared according to the protocol in the previous study^39^. For HDR-mediated knock-in, 1.25 µl of 20 µM EnGen Spy Cas9 NLS protein (New England Biolabs) was combined with 2.0 µl of 12 µM gRNA, 0.5 µl of 500 ng/µl Alt-R single-stranded donor DNA, and 0.5 µl of 0.5% phenol red, and the volume was adjusted to 5 µl with 2 M potassium chloride. Chemically modified single-stranded donor DNAs were obtained from Integrated DNA Technologies. A mixture of Prime Editor or Cas9 protein and gRNA was incubated at 37°C for 5 minutes prior to microinjection to form an RNP complex. Glass needles for microinjection were prepared from glass capillaries (TW100F-4, World Precision Instruments) using a P-1000 micropipette puller (Sutter Instrument). Approximately 1–1.7 nl of the injection mixture was injected per embryo. After microinjection, the injected embryos were incubated in E3 embryo medium (5 mM sodium chloride, 0.17 mM potassium chloride, 0.33 mM calcium chloride, and 0.4 mM magnesium chloride; pH 7.2) at 32°C, following the protocol by Petri et al^18^.

### DNA extraction, PCR analyses and molecular cloning

Genomic DNA was extracted from individual embryos and larvae using the modified HotSHOT method^74^. The target region for prime editing in *ror2*, *adgrf3b*, and *smyhc1:gfp* was amplified by PCR using the following primer sets: *ror2* (5’-AAACTTATGGGTGCCAGTCC-3’ and 5’-ATGGACACAAACTGAGGCTG-3’), *adgrf3b* (5’-TGATTGCATACACACCTGACC-3’ and 5’-AGGCACCTGCAGGAAAATTA-3’), and *smyhc1:gfp*(5’-TGCAGTTACAAGGTACAGAGGTC-3’ and 5’-CGTCCTTGAAGAAGATGGTGCG-3’). The PCR mixture was prepared as follows: 10 µl of 2x PCRBIO Taq Mix red (PCR Biosystems), 0.8 µl each of 10 µM forward and reverse primers, 1 µl of genomic DNA pooled from 10 individual samples, and 7.4 µl of water, totalling a volume of 20 µl. The PCR settings were as follows: 95°C for 1 min; 28 cycles of 95°C for 15 s, 58°C for 15 s, and 72°C for 15 s, followed by 72°C for 1 min. The mutation in the target sequence of the PCR product was assessed by the T7 endonuclease I (T7E1) assay. PCR products were denatured at 95°C for 5 min and then cooled to room temperature to form heteroduplex DNAs. Subsequently, 5 µl of PCR product was mixed with 1 µl of NEBuffer 2 (New England Biolabs), 0.15 µl of T7E1 enzyme (10 U/µl, New England Biolabs), and 3.85 µl of water to a total volume of 10 µl and incubated at 37°C for 15 min. The PCR and T7E1-digested products were assessed on a 3% agarose gel alongside a 100 bp DNA ladder (New England Biolabs). PCR products were also cloned into the pGEM-T Easy vector (Promega) and used for transformation with One Shot TOP10 chemically competent cells (Invitrogen) following the manufacturer’s instructions. Bacterial colonies grown on an LB agar plate with 100 µg/ml ampicillin were randomly selected and used for plasmid extraction using a GeneJET plasmid miniprep kit (Thermo Scientific). Sanger sequencing of PCR products and plasmids was performed by Eurofins Genomics.

### Amplicon sequencing and bioinformatics

PCR amplicons for next-generation sequencing (NGS) of *adgrf3b*, *crbn*, *ror2*, *wls*, and *smyhc1:gfp* were prepared using Phusion Flash High-Fidelity 2x Mastermix (F548, Thermo Scientific) in a reaction volume of 15 μl, which contained 1.5 μl of genomic DNA extract and 0.2 μM of specific primers with barcodes and adapters for NGS. All primer sequences are shown in Supplementary Table 2. The PCR protocol with Phusion Flash High-Fidelity 2x Mastermix included an initial step at 98°C for 3 min, followed by 30 cycles at 98°C for 10 s, 60°C for 5 s, and 72°C for 5 s. The resulting PCR amplicons were purified using the HighPrep PCR Clean-up System (MagBio Genomics). The characterisations of size, purity, and concentration of the amplicons were conducted using a fragment analyser (Agilent). A second PCR was carried out to add Illumina indexes to the amplicons, utilising KAPA HiFi HotStart Ready Mix (Roche) in a total volume of 25 µl, containing 0.067 ng of PCR template and 0.5 µM indexed primers (Illumina). The PCR conditions were set at 72°C for 3 min, 98°C for 30 s, followed by 10 cycles at 98°C for 10 s, 63°C for 30 s, and 72°C for 3 min, with a 5-minute final extension at 72°C. Amplicons were purified with the HighPrep PCR Clean-up System (MagBio Genomics) and analysed using a fragment analyser (Agilent). Quantification was performed using a Qubit 4 Fluorometer (Life Technologies), with sequencing carried out on the Illumina NextSeq system in accordance with the manufacturer’s guidelines. Demultiplexing of the amplicon sequencing data was executed with bcl2fastq software. The resulting fastq files were processed with CRISPResso2 V2.2.12 in prime editing mode^75^. Detailed parameters of the CRISPResso analysis can be found in Supplementary Table 3.

### Live imaging

Stereomicroscope images were obtained using an Olympus DP73 microscope camera, and measurements were conducted with cellSens imaging software (Olympus). Confocal images of *smyhc1:gfp* transgene expression were captured with a Leica TCS SP8 laser scanning microscope using a HC Fluotar L 25×/0.95 water objective and LAS X imaging software (Leica). For imaging, live zebrafish larvae were affixed to glass slides with 1% low melting agarose (5806A, Takara) in an E3 embryo medium containing 200 µg/ml tricaine (A5040, Sigma).

### Computed Tomography

Adult wild-type and *ror2*^W722X^ mutant zebrafish at 12 months were fixed in 4% PFA for one week and dehydrated in 70% ethanol. Full body scans were taken using a Nikon XT H 225ST CT scanner (X-ray source of 130 kV and 53 μA without additional filters) with a voxel size of 20 μm. The generated scans were reconstructed using CT Pro 3D software (Nikon). During reconstruction, greyscale values were calibrated against a scan of a phantom with known density (0.75 g/cm^3^). To evaluate shape changes while controlling for isometric growth, ratios of linear distances between annotated geographical points in three-dimensionally segmented reconstructions were calculated. Image renders and morphological analyses were performed using volume rendering function of Avizo software (FEI).

### Statistics

Data visualisation and statistical analyses were conducted using GraphPad Prism 10.4 (GraphPad Software), BioRender Graph (R version 4.2.2), and Adobe Illustrator. For statistical analysis, the numerical dataset from each experimental condition was examined using the Shapiro-Wilk test for normality and Levene’s test for homogeneity of variance. One-way ANOVA with Tukey multiple comparison test was utilised to assess differences between three or more groups with similar standard deviations, and Welch’s one-way ANOVA with Dunnett T3 multiple comparison test for the groups having unequal variances (parametric). The Kruskal-Wallis test with Dunn’s multiple comparison test was utilised to assess differences between the groups that are not normally distributed (nonparametric). Unpaired t-test (parametric) and the Mann-Whitney U test (nonparametric) were employed to evaluate differences between the two groups. The sample size for each experiment is presented in the figures and legends.

## Acknowledgements

Research in the S.S. lab, including Y.O. and A.B., is supported by the Biotechnology and Biological Sciences Research Council Industrial Partnership Awards with AstraZeneca (BBSRC-IPA, BB/X008401/1; awarded to S.S., Y.O., and C.R.T) and the Living Systems Institute Open Innovation Platform Fund at the University of Exeter (awarded to Y.O.). M.L. receives support from the National Centre for the Replacement, Refinement and Reduction of Animals in Research (NC3Rs) PhD studentship (NC/X001407/1; awarded to J.S.B and C.R.T). C.L.H acknowledges funding from Versus Arthritis Senior fellowship (29317). A.K. was funded by GW4 BioMed MRC Doctoral Training Partnership, and F.B. by the South West Biosciences Doctoral Training Partnership (SWBio-DTP). We thank the Aquatic Resources Centre at the University of Exeter for the care of zebrafish resources.

## Author contributions

Y.O., M.P., M.M., and S.S. conceived the study and designed the experimental strategy. Y.O., M.L., and A.B. performed the animal experiments and Y.O. and M.P. analysed the data with support from E.G., J.S.B., C.R.T., S.R., and M.B. Computed Tomography and the image analysis were conducted by A.K., F.B., and C.L.H. The manuscript was prepared by Y.O. and S.S.

## Competing interests

M.P., E.G., S.R., M.B., and M.M. are employees of AstraZeneca and may be AstraZeneca shareholders.

## Notes

### Competing Interest Statement

The authors have declared no competing interest.

### Summary of Updates

In the revised version, we - Added the late phenotype analysis of the ror2 mutant, including loss of nasal and maxillary barbels and altered adult jaw morphology by microCT, strengthening the disease-model relevance. The microCT analysis was performed in collaboration with the Hammond lab; therefore, Amir Khan, Felix Bowers, and Chrissy L. Hammond were added as co-authors. - Added new data on a further target locus (wls) showing 46 bp attP insertion by PEn and comparison with HDR-mediated knock-in at the same site. - Expanded the analysis of insertion performance at adgrf3b and clarified comparison with previously reported PE2 data. - Added the analysis of HDR-mediated knock-in and prime editing substitution to generate ror2 W722X allele. - Added comparative off-target analysis for PE2, PEn and HDR at three predicted off-target sites for the ror2 target. - Resolved the cloning/NGS inconsistency for ror2 by increasing clone analysis

