## Supplementary material for "Optimised genome editing for precise DNA insertion and substitution using Prime Editors in zebrafish": Suppl data

### Supplementary Figures

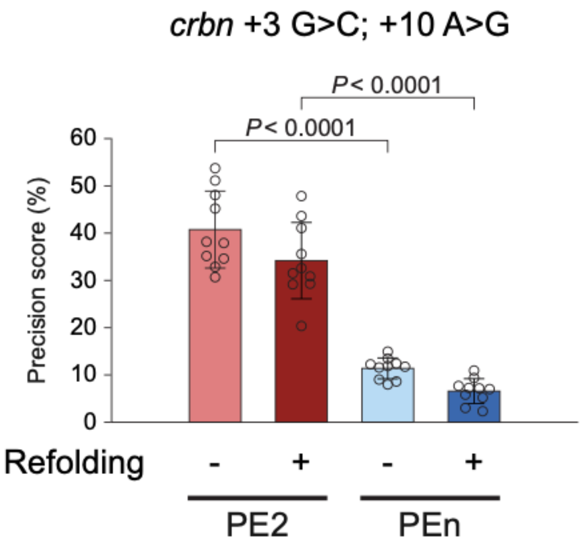


**Supplementary Fig. 1 Prime editing substitution in the zebrafish *crbn* gene.** Quantitative analyses of precision scores comparing between PEn and PE2 in prime editing substitution in the *crbn* gene with the pegRNA refolding procedure (n = 10 per group). P-values were determined using Welch's one-way ANOVA with Dunnett T3 multiple comparison test. Error bars in the bar graphs represent the mean and standard deviation, and individual data points indicate values from single injected embryos.

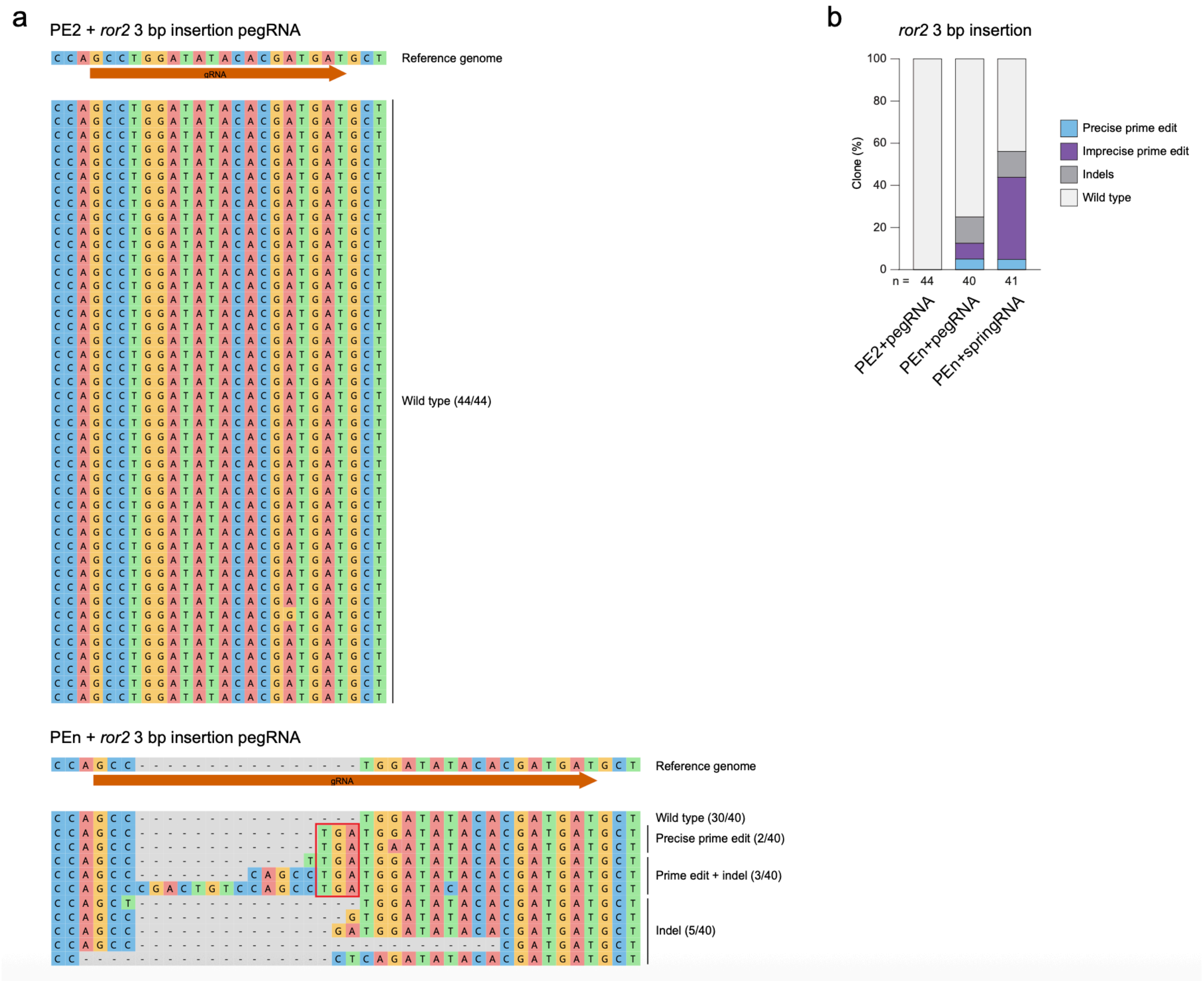


**Supplementary Fig. 2 Analysis of prime editing insertion in zebrafish *ror2* gene by sequencing the clones.** a, Sequence alignment of the edits in the *ror2* target site obtained from randomly selected bacterial clones comparing prime editing outcomes using PE2/pegRNA (top) and PEn/pegRNA (bottom) combinations. The guide RNA (gRNA) target sequence is indicated by red arrows. b, Quantitative comparison of editing outcomes using different combinations of Prime Editor and gRNA. The proportion of the clones with each type of edit is presented.

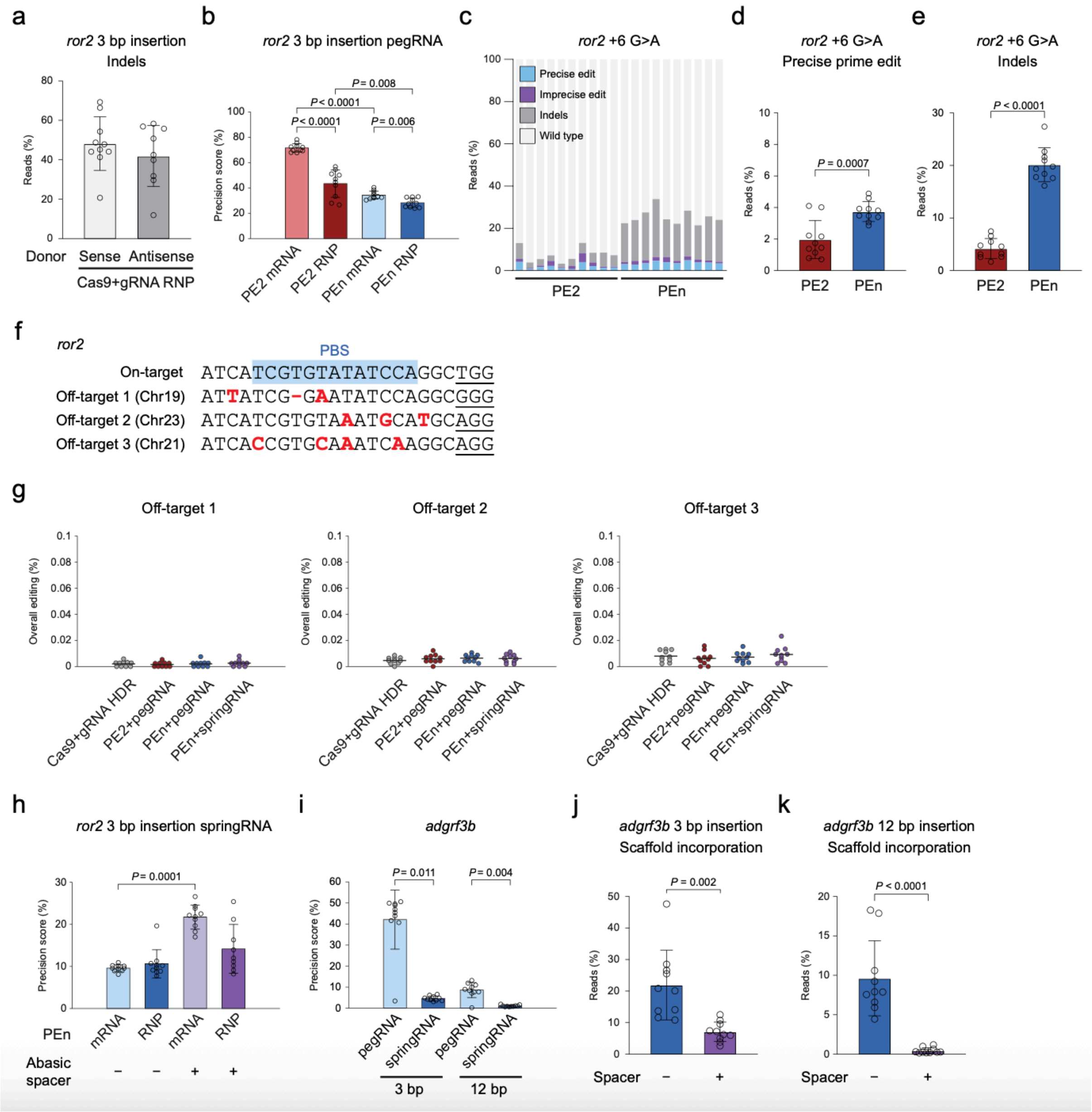


**Supplementary Fig. 3 Amplicon sequencing analysis of prime editing insertion in zebrafish *ror2* and *agdrf3b* genes.** a, Proportion of reads with indels in HDR-mediated knock-in in *ror2* using single-stranded sense or antisense donor DNAs with 40 bp homology arms. b, Quantitative analyses of precision scores comparing various experimental conditions for prime editing insertion in *ror2* gene using pegRNA. c–e, Comparison between PEn and PE2 in prime editing substitution in *ror2* to generate the W722X allele. The proportion of editing outcomes in individual injected embryos was assessed through amplicon sequencing (c), along with a quantitative analysis of precise prime edit (d) and indels (e) across experimental conditions. f, Comparison of potential off-target sequences of *ror2* pegRNA and springRNA. Mismatched bases and a gap are indicated in bold red and primer binding site (PBS) is highlighted in blue.

PAM sequences are underlined. g, Proportion of overall editing in each off-target site comparing HDR-mediated knock-in and various prime editing conditions. h and i, Precision scores in prime editing insertion in *ror2* gene using springRNA (h), and in *adgrf3b* gene using PEn mRNA (i). j and k, Proportion of reads with scaffold incorporation in 3 bp (j) and 12 bp (k) prime editing insertion in *adgrf3b* using springRNAs with an RNA (j) or DNA (k) spacer. The sample size is n = 10 per group and one sample of the PEn RNP/abasic springRNA combination was excluded from analysis in h due to low read count. P-values were determined by Welch's one-way ANOVA with Dunnett T3 multiple comparison test in b, and Kruskal-Wallis test with Dunn's multiple comparison test in h and i. To evaluate differences between the two groups P-values were determined by unpaired t-test in d and e. Welch's t-test and the Mann-Whitney U test were employed in j and k, respectively. Error bars in bar graphs represent the mean and standard deviation, and each individual data point indicates the value from a single injected embryo.

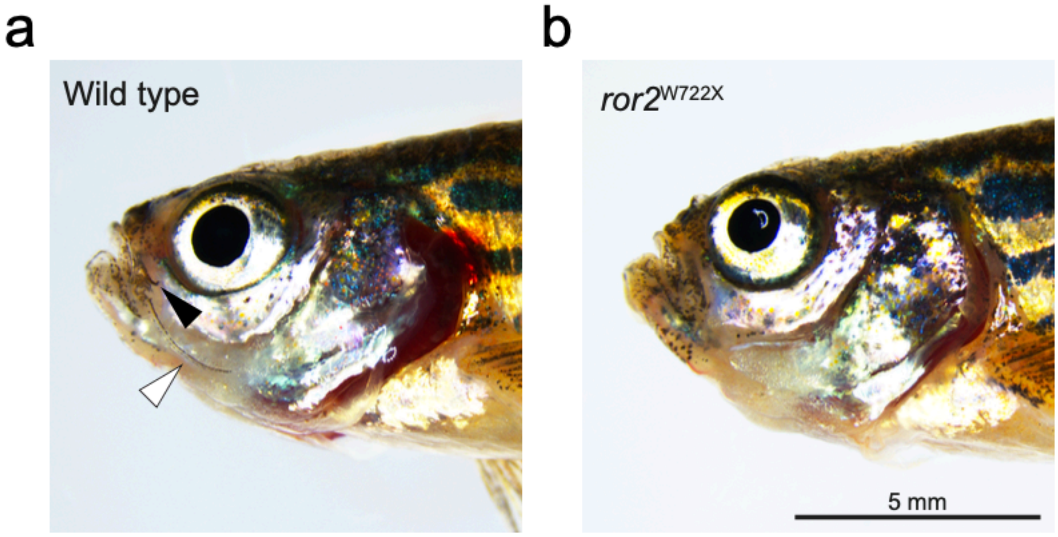


**Supplementary Fig. 4 Characterisation of stable *ror2*$^{W722X}$ mutant.** a and b, Lateral images of 18 months old wild type (a) and zygotic *ror2*$^{W722X}$ mutant (b). Nasal and maxillary barbels are indicated by black and white arrowheads, respectively.

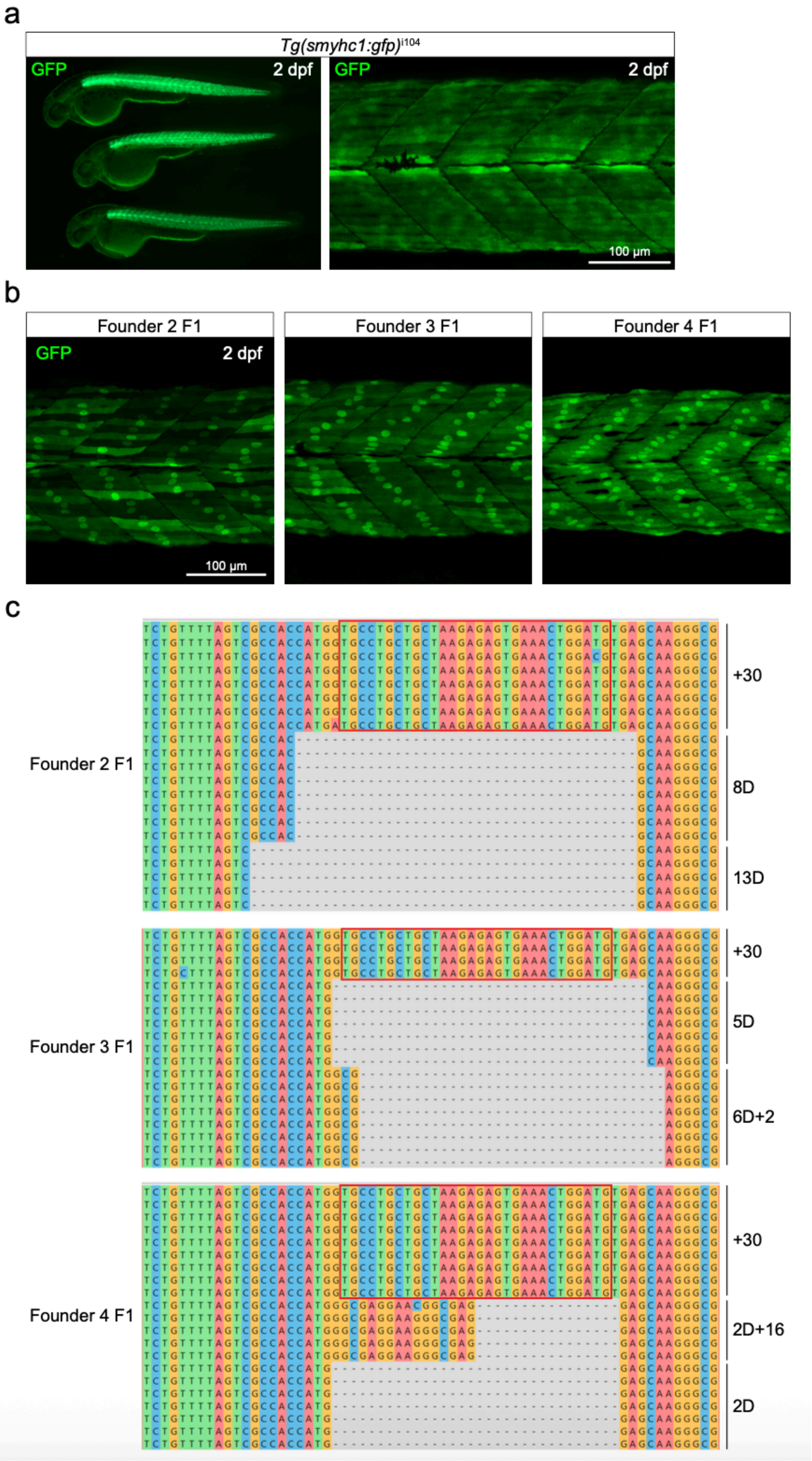
a
Tg(smyhc1:gfp)i104
GFP
2 dpf
100 µm
b
Founder 2 F1
Founder 3 F1
Founder 4 F1
c
+30
8D
13D
5D
6D+2
2D+16
2D

**Supplementary Fig. 5 Prime editing insertion of a nuclear localisation signal sequence into the *smyhc1:gfp* transgene.** a, The expression of *smyhc1:gfp* transgene in 2 days post-fertilisation (dpf) larvae. Lateral images of whole larvae (left) and the trunk muscle (right). Anterior is to the left. b, Confocal microscopy images of the trunk muscle of F1 larvae exhibiting nuclear GFP expression at 2 dpf, obtained from founder 2, 3, and 4 (see Fig. 5f). c, Sequence alignment of the edits in the target site obtained from F1 embryos exhibiting nuclear GFP expression (founder 2, 3, and 4). The nuclear localisation signal (NLS) sequences are outlined in red.

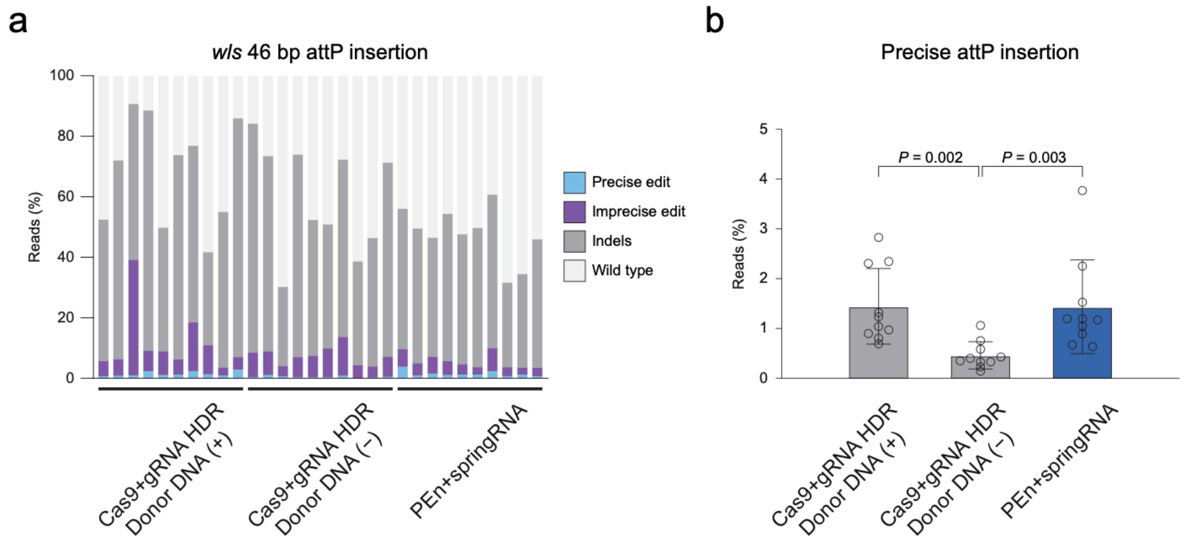


**Supplementary Fig. 6 Prime editing insertion of a 46 bp attP sequence into zebrafish *wls* gene.** a and b, Comparison between HDR-mediated CRISPR/Cas9 knock-in and prime editing using PEn/springRNA RNP complex in inserting 46 bp attP sequence into *wls* gene. Two single-stranded donor DNAs with 40 bp homology arms in the same (+) or opposite (−) directions of the target spacer sequence of guide RNA were used. The proportion of editing outcomes in individual injected embryos was assessed through amplicon sequencing (a), along with a quantitative analysis of precise edit (b) across experimental conditions (n = 10 per group). P-values were determined using the Kruskal-Wallis test with Dunn's multiple comparison test. Error bars in the bar graphs represent the mean and standard deviation, and individual data points indicate values from single injected embryos.
